# Bicoid gradient formation mechanism and dynamics revealed by protein lifetime analysis

**DOI:** 10.1101/280834

**Authors:** Lucia Durrieu, Daniel Kirrmaier, Tatjana Schneidt, Ilia Kats, Sarada Raghavan, Michael Knop, Timothy E Saunders, Lars Hufnagel

## Abstract

Embryogenesis relies on instructions provided by spatially organized signaling molecules known as morphogens. Understanding the principles behind morphogen distribution and how cells interpret locally this information remains a major challenge in developmental biology. Here we introduce morphogen-age measurements as a novel approach to retrieve key parameters in morphogen dynamics. Using a tandem fluorescent timer (tFT) as a protein-age sensor we find a gradient of increasing age of Bicoid (Bcd) along the anterior-posterior (AP) axis in the early *Drosophila* embryo. Quantitative analysis retrieves parameter that are most consistent with the synthesis-diffusion-degradation (SDD) model underlying Bcd-gradient formation, and rule out some other hypotheses for gradient formation. Moreover, we show that the timer can detect transitions in the dynamics associated with syncytial cellularization. Our results provide new insight into Bcd gradient formation, and demonstrate how morphogen age-information can complement knowledge about movement, abundance and distribution, which should be widely applicable for other systems.

---

Acquisition of different cell fates at specific spatial and temporal locations is an essential process driving development. The necessary information is provided locally by morphogens^1,2^. To achieve this, morphogen expression and distribution across tissues is controlled precisely in a spatially confined manner. To understand morphogen gradient formation requires systematic measurement of the morphogen abundance, mobility and distribution using temporally resolved methods. The technical challenges associated with this undertaking are yet unsolved, with significant discussion on how to best assess the principles and mechanisms of their transport^3-5^. This ambiguity in the data and its interpretation has resulted in a plethora of models for formation of morphogen gradients.

In the early fly embryo, the morphogen Bicoid (Bcd) forms a gradient with high concentration in the anterior pole that determines cell fate along the axis to the posterior pole in a concentration dependent manner^6,7^ (Fig. 1a). The process is initiated during oogenesis where Bcd mRNA (*bcd*) is localized to the anterior of the forming embryo^3-5,8,9^. The classic view of Bcd gradient formation is that the protein is synthesized in the anterior pole of the *Drosophila* blastoderm and forms a long range gradient through diffusion, with the gradient shape adapted by protein degradation (SDD model)^6,7,10^. Such a model fits well with the observed Bcd gradient in nuclear division cycle 14 (n.c.14), where Bcd levels decay exponentially towards the posterior pole^10,11^. However, several other models involving alternative mechanisms for Bcd production and distribution have been proposed, all of which are capable of producing an exponential-like concentration profile, as further outlined below^12-17^.

**Figure 1.**
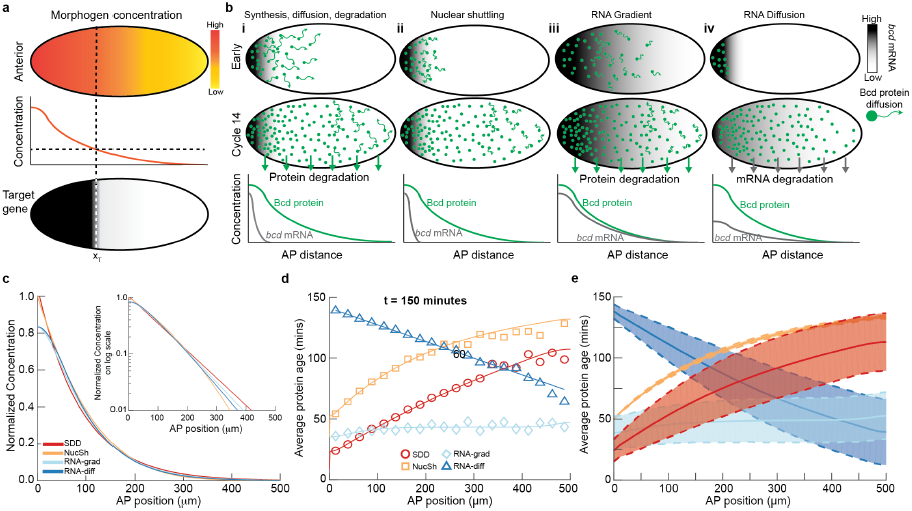
Protein age can distinguish alternative models of morphogen gradient formation with similar concentration profiles. (a) Cartoon of the morphogen hypothesis: a spatially varying concentration of signaling molecule can result in precise readout of positional information. (b) Outline of models considered.Morphogen RNA (grays) and protein (green dots) distribution are shown in early (top) and cycle 14 (middle) rows for each model considered. The magnitude of protein movement is represented by length of green arrows. Degradation of protein/RNA is represented by green/gray arrows. Bottom row shows a schematic of morphogen RNA and protein concentration profiles in cycle 14. (c) Normalized Bcd concentration profiles for the models considered in b at time=2.5 hours. Inset: same on log scale. See Supplementary Information and Supplementary Fig. 1 for extended discussion of models and parameters. (d) Simulation results for average protein age as function of position, 2.5 hours after initiation. Colored lines correspond to theoretical predictions for protein age in each model. (e) As d but showing solutions over larger parameter space. Solid line represents mean solution and dashed lines represent 1s.d. Parameter range described in text. SDD: synthesis, diffusion, degradation model, NucSh: nuclear shuttling model, RNA-grad: RNA gradient model, and RNA-diff: RNA diffusion model.

Efforts to distinguish the different mechanisms have been hindered by uncertainties associated with the measurements of the parameters needed to quantitatively validate different models: local production rates of Bcd; Bcd mobility and transport; and Bcd degradation. Experimental estimates of the diffusion constant vary by an order of magnitude^10,18^, although recent theoretical work has attempted to meld these measurements^19,20^, while *in vivo*^21^ and *in vitro*^22,23^ measurements of the Bcd degradation rate also significantly differ. These differences in the diffusion and degradation rates are meaningful, as they help determine whether the system is in (or close to) equilibrium – a contested issue relevant to the mechanism of gradient interpretation^24-27,28-30^ - or indeed whether diffusion alone is sufficient to form the gradient profile^12^. Finally, *bcd* mRNA is not strictly localized to the anterior pole^13,31^, suggesting spatially distributed Bcd protein production. Altogether, these debates regarding nearly every aspect of Bcd gradient formation argue for the need of more incisive tools to investigate this paradigmatic problem.

In this work, we revisit Bcd gradient formation based of the observation that the measured protein gradient changed shape as a function of different fluorescent proteins fused to Bcd^31,32^. Differences in the fluorophores maturation rates could underlie this change, as discussed^31,32^. Here we employ the tandem fluorescent protein timer (tFT) reporter^33^, which provides simultaneously quantitative information about the Bcd protein age and its spatio-temporal concentration distribution. We use multi-view light-sheet fluorescent microscopy^34,35^ to gain high spatial and temporal resolution images of the tFT-Bcd *in toto*. This data is then used to discriminate between models of Bcd gradient formation, estimate dynamic parameters, investigate the effect of a mutation in a gene previously implicated in Bcd degradation, and study temporal changes through early fly embryogenesis.

## Results

### Protein age can distinguish different models of Bcd gradient formation

In Bcd gradient formation, broadly two types of models have been considered: localized Bcd synthesis in the anterior with subsequent long-ranged transport; and pre-patterned synthesis (by an RNA gradient) and restricted protein transport, although also more complicated scenarios such as spatially patterned degradation are imaginable. Interestingly, protein degradation is not a mandatory ingredient in either scenario but plays an important role in determining whether the system can reach steady-state. Within this framework, we consider four models: the SDD model^6,10^; the *nuclear shuttling model*^12^; *bcd mRNA gradient* (ARTS)^13,17,36^; and *bcd mRNA diffusion and degradation*^15,36^. In the SDD and nuclear shuttling models, protein is synthesized locally and then migrates by diffusion towards the posterior pole (Fig.1b,i-ii). The SDD model incorporates protein degradation, whereas the nuclear shuttling model utilizes the rapid increase in nuclei number in the blastoderm to enable the Bcd concentration to remain roughly constant in each nucleus. The RNA gradient model (Fig. 1b, iii) is based on an RNA gradient under-laying distributed protein synthesis and incorporates protein degradation and very slow protein diffusion. The RNA diffusion model starts with localized RNA and protein synthesis, and then proposes spreading of the mRNA (protein synthesis) throughout the embryo (Fig. 1b, iv). This model reaches a steady-state when the RNA is completely degraded, even though Bcd protein does not decay (and the Bcd protein cannot diffuse)^15^. Further details about the models are provided in Supplementary Information.

All four models can reproduce the observed Bcd concentration profile at n.c.14, as expected (Fig. 1c, Supplementary Fig. 1a-b). Distinguishing these models experimentally requires precise measurement of dynamic parameters – which has proven challenging so far. Thus, easily measurable information is required where the models make distinct and robust predictions. What information could this be?

It has been shown that tandem fusions of two different fluorescent proteins with different maturation rates can measure the average time that has passed since the production of a pool of proteins (*i.e.* its age)^33^. Protein age is dependent on protein turn-over and degradation. Thus, we reasoned that fluorescent timers could be a valuable tool for discerning these models experimentally, which prompted us to explored their predictions regarding protein age. In models that include degradation, the average age of Bcd approaches a steady-state situation where synthesis and degradation are balanced. In models without protein degradation, the average protein age is constantly increasing (Supplementary Fig. 1c). We calculated the Bcd protein age as a function of position in the AP axis (Fig. 1d, Supplementary Fig. 1c-d and Supplementary Information). For the SDD model and the nuclear shuttling models, protein age increases with the distance from the anterior pole, though the average protein age in the SDD model is shorter due to degradation. For the RNA gradient model - where the *bcd* RNA gradient is the primary determinant of the Bcd gradient - the average age of Bcd is roughly uniform. In contrast, the RNA diffusion model makes an inverse prediction, with younger protein age towards the posterior pole.

To test the robustness of these predictions we explored the parameter space underlying each model for the situation after 2.5 hours (around n.c.14). We varied each parameter over a physiologically relevant range: diffusion coefficient (0.1 to 10 μm^2^s^-1^), protein and RNA lifetimes (10-120 minutes) and the range of the RNA gradient (20-200 μm). We restricted the gradients to exponential-like profiles with decay lengths between 70-100 μm. This revealed that the principal difference of the models with respect to relative Bcd protein age in the gradient is robust, and should allow faithful discrimination between the models (Fig 1e and Supplementary Fig. 1e). We note that distinguishing the SDD and nuclear shuttling models is the most challenging. However, combining the protein age data with the protein concentration profiles should enable to discriminate between the two models.

### Establishment of a protein-age reporter line

To quantify Bcd protein age experimentally, we fused N-terminally tFTs composed of super-folder GFP (sfGFP^37^) and different red fluorescent proteins, and expressed them in embryos keeping the regulatory 5’ and 3’ sequences of the *bcd* gene (Fig. 2a). A pool of proteins tagged with the tFT tag changes its color as a function of time (Fig.2b) with a dynamic range that mostly depends on the slower (red) fluorescent protein. Moreover, to be informative, the timescale for maturation of the slower maturing fluorophore must be not slower than the dynamics of the system studied (in our case, on the order of an hour)^33,38^. We constructed six tFT-Bcd lines, each containing different red FPs and analyzed the red and green signal intensity along the AP axis. Based on this screening, we chose two lines for further investigation (Supplementary Fig. 2): a line tFT-Bcd (consisting of sfGFP and mCherry tagged to Bcd), and one containing a faster maturing mCherry that we developed using directed protein evolution, termed fmCherry (Methods). We also established a control line that expresses only the tFT reporter (without the Bcd protein coding sequence), but using the regulatory sequences of the *bcd* gene. We validated the functionality of the tFT-Bcd fusions by rescuing the *bcd*^*E1*^/*bcd*^*E1*^ null mutant^6^ (Supplementary Fig. 3a-b).

**Figure 2.**
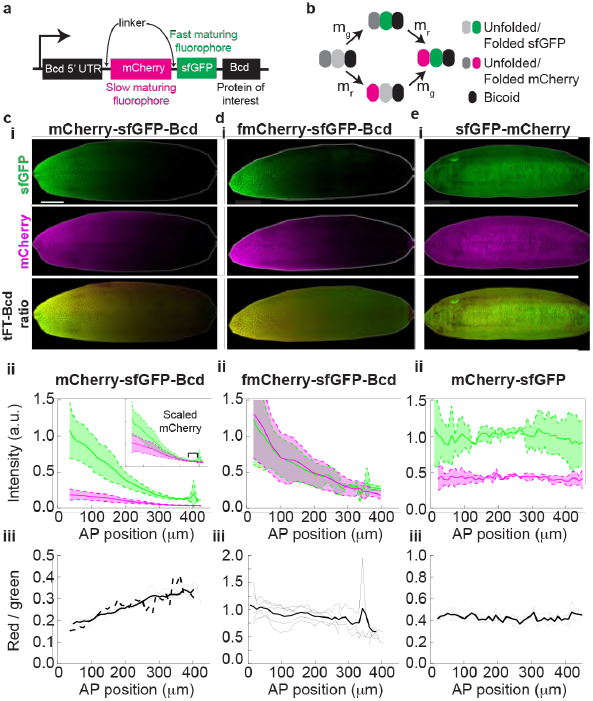
tFT-Bcd reporter reflects average protein age. (**a**) Schematic of tFT-Bcd reporter with mCherry and sfGFP fluorophores. (**b**) Cartoon of the fluorescent protein maturation states in the tFT-Bcd reporter. m_g_ and m_r_ represent the sfGFP and mCherry maturation rates respectively (note, m_r_ is an effective rate as mCherry has a two-step maturation process). (**c**,**d**,**e**) Examples of embryos expressing the tFT-Bcd reporter with **c**: mCherry-sfGFP-Bcd, **d**: fmCherry-sfGFP-Bcd, and **e**: mCherry-sfGFP. For all, (**i**) Images of shells (see Methods) of embryos in early n.c. 14. (**ii**) Mean AP intensity profile of each color for the embryo in **i**. Shade region represents ± 1 s.d. Inset shows same profiles after multiplication of red intensity by constant factor. Data binned into 10 μm bins (n=4 embryos). (**iii**) Ratio of green over red signal, reflecting protein age. The thin lines represent individual embryos, while the thick solid line is the mean. The solid dashed line depicts the mean green/red ratio for a line with the tFT-Bcd reporter but lacking endogenous Bcd (Bcd^E1^ mutant, n=4, single embryos not shown).

**Figure 3.**
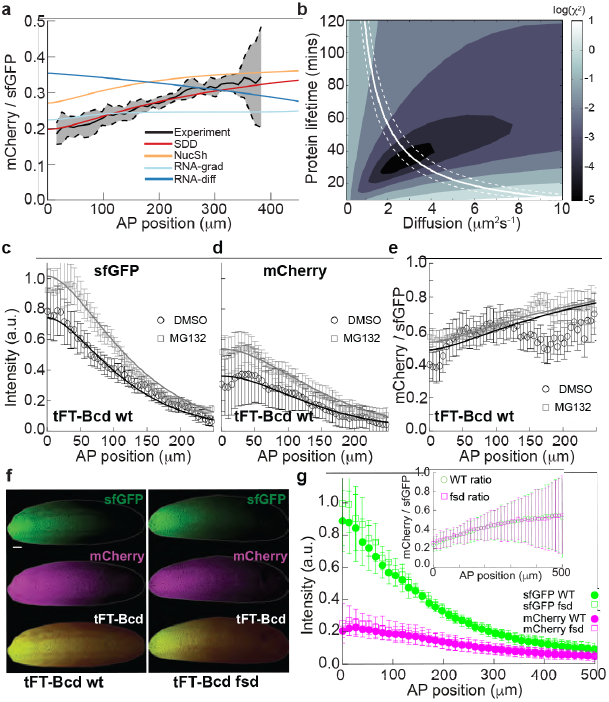
Estimation of Bcd diffusion and degradation rates supports the SDD model and Fsd mutation does not significantly alter Bcd lifetime. (**a**) The SSD model fits best the timer reporter experimental data. Model fitting to the experimental tFT-Bcd ratio in early n.c. 14 as function of AP position for the different models outlined in Fig. 1a. Black line represents mean measured ratio of tFT-Bcd reporter with mCherry-sfGFP-Bcd in y/w flies. Dashed lines represent ± 1s.d. (**b**) Fitting of the timer reporter data allows independent determination of diffusion and degradation rates. Quality of SDD model fitting to tFT-Bcd ratio (showed in **a**). The colormap is the log of the fitting function (Methods). The estimated values for the parameters are: D = 3.0 ± 0.7 μm^2^s^-1^, t_1/2_= 38 ± 7 minutes. The solid (dashed) white line corresponds to parameter combinations that produce a gradient with a length constant *λ* = 82.5 *μm* (*λ* = 90 *μm* and *λ* = 0075 *μm*). (**c**,**d**,**e**) Bcd is degraded by the proteasomes *in vivo*. Injection of the proteasome inhibitor MG132 embryos increases both the amount and age of Bcd (gray, open squares, N=10). Control embryos were injected with DMSO (black, filled circles, N=9). All embryos are y/w and were injected while in cycle 14 and imaged 60 min later. (**f**) The age of Bcd and its gradient are unaltered in *fsd* mutant. Comparison of a representative y/w embryo (**left**) and loss-of-function mutation fsd^KG02393^ (**right**) are shown. Both embryos have the tFT-Bcd construct in the same locus, are in early n.c. 14, were imaged side-by-side under identical conditions and are displayed with the same settings. (**g**) Quantification of embryos as in **f**. Fluorescence intensity from sfGFP (green) and mCherry (magenta) for wt (N=8, filled circles) and fsd (N=6, open squares) at early cycle 14. **Inset**: tFT-Bcd ratio for wt (black) and FSD (gray) embryos.

### Quantification of the tFT-Bcd in embryos

Reliable quantification of fluorescence signals in the *Drosophila* embryo *in toto* is challenging. For this we used a confocal multi-view light-sheet microscope^34,39,40^. Light sheet-imaging enables highly sensitive *in toto* fluorescence detection in larger specimens (e.g. embryos), while bleaching and phototoxicity is strongly reduced compared to confocal detection methods^41^. For quantification of tFT-Bcd signals, we first reduced the 3D data to 2D by a mapping procedure that generates ‘ carpets’ ^34^ that quantitatively represent the area of the embryonic periphery where the nuclei reside (Supplementary Fig. 4a and b; Methods). Validation of our methods, including the microscope sensitivity can be found in Supplementary Fig. 4c-f, and Supplementary Information. We can detect Bcd-GFP fluorescence signal as early as n.c.8, (Supplementary Fig. 4g), and in n.c.14 we detected similar intensity variability to previously reported fluctuations in Bcd^10^ and above-background Bcd signal in all nuclei, even at the posterior pole, Supplementary Fig. 4h-l.

**Figure 4.**
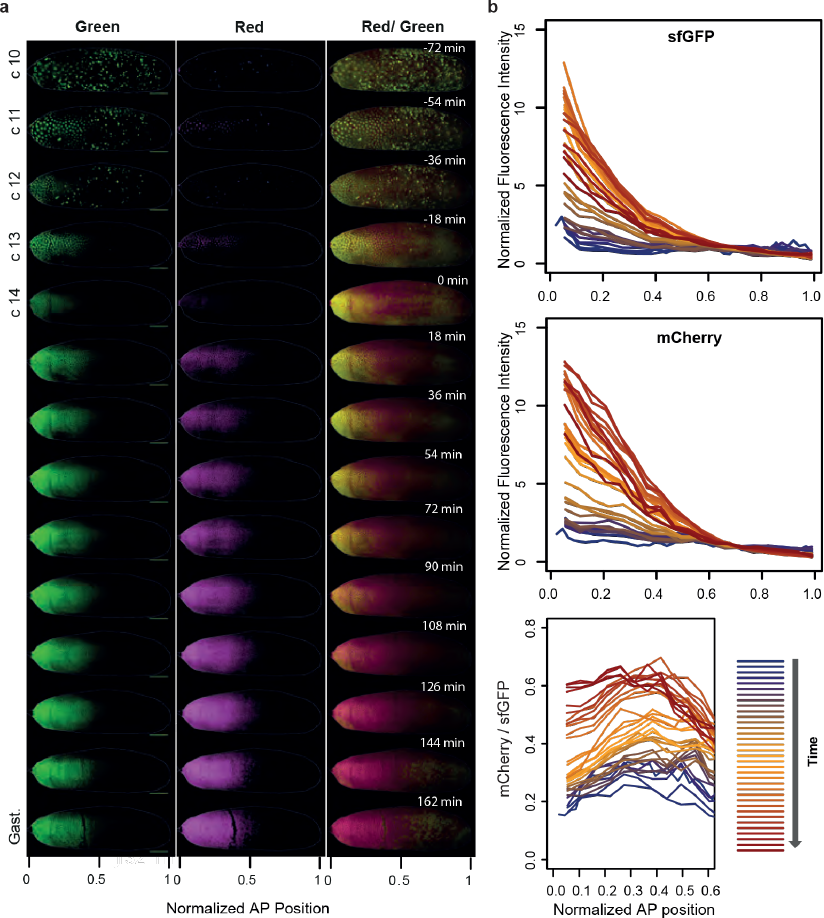
The average age of Bcd proteins increases continuously during the cleavage divisions and the gastrulation. Time-course of a representative y/w embryo with the tFT-Bcd reporter from the division cycle 10 to the early gastrulation. A movie of the same embryo with 6 min time resolution is available as Supplementary Video 1. (**a**) SPIM images of the embryo shell (see methods). **Left**: Green fluorescence, **center**: Red fluorescence, **right**: Weighted ratiometric image (see Methods) of the mCherry/sfGFP ratio, reporting average protein age. (**b**) Quantification of the same embryo as is **a**.

Quantification of the tFT-Bcd fluoescence- in early n.c.14 embryos revealed that the mCherry/sfGFP intensity ratio increased along the AP axis (Fig. 2c). This implies that, on average, the tFT-Bcd is older towards the posterior. In contrast to this, the fmCherry-sfGFP-Bcd line showed a weak but reproducible reversed ratio behavior, Fig. 2d. fmCherry was evolved in yeast based on selection for increased maturation rates (Methods). The most likely explanation would be that fmCherry is maturing with similar speeds or even slightly faster than sfGFP. To test this, we quantified fluorophore formation rates and obtained for sfGFP and fmCherry maturation rates in the range of 20 min, whereas mCherry maturation was around 50 min (Supplementary Fig. 5a-c, Supplementary Information). To check that the observed tFT ratio reflects the age of the Bcd protein and not itself, we tested the line with the mCherry-sfGFP reporter without the Bcd protein. This revealed no intensity or tFT ratio gradients, Fig. 2e. A known possible artifact of tFT measurements due to incomplete degradation of the fluorescent timer was excluded^33,42^, (Supplementary Fig. 5b, Supplementary Information). In conclusion, our quantitative analysis demonstrates an increase in the relative Bcd age toward the posterior pole and this result does not depend on the specific scaling of the measured intensities (Supplementary Fig. 2m-n). This is qualitatively in agreement with the predictions from the SDD and nuclear shuttling models (Fig. 1d).

**Figure 5.**
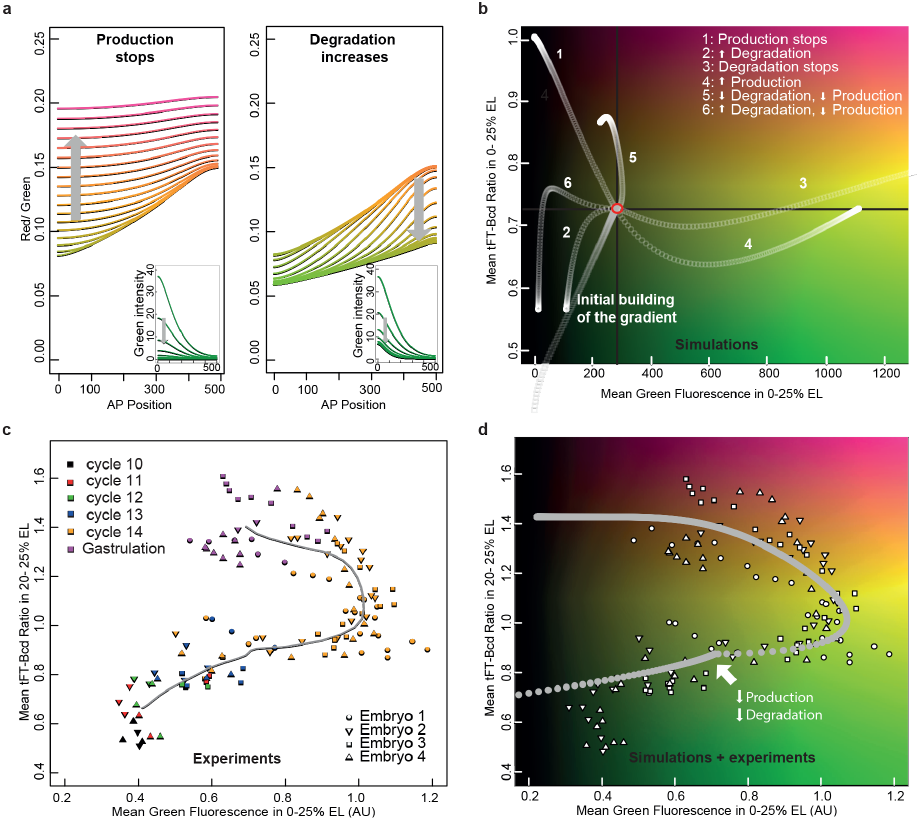
At the onset of cellularization Bcd production and degradation rates decrease. (**a**) Simulation of SDD model, where after reaching steady-state (orange line), either production was stopped (**left**) or degradation increased (**right**). The gray arrow shows the direction of time, the color of the lines reflects the tFT ratio (y axis).(**b**) As **a**, where the SDD model was run until reaching steady state (gray circles, red circle) and then the following perturbations were performed (1) Degradation was stopped, (2) Production rate was increased 2-fold, (3) Degradation rate was increased 2-fold, (4) Synthesis was stopped, (5) synthesis and degradation were decreased 2-fold, (6) production decreases and degradation were increased 2-fold. The y-axis shows the mean red/green ratio (proxy for average protein age) in the anterior region of the embryo (0 to 25% of embryo length) and the x-axis represents the mean green fluorescence intensity in the same region (age/intensity diagram). (**c**) During the cleavage divisions the embryos define reproducible trajectories in the age/intensity diagram. Experimental data from quantification of movies of y/w embryos as in Fig.4. The trajectories of 4 embryos from n.c. 10 to gastrulation are shown (each embryo depicted with a different symbol). The data points are colored according to the division cycle. The intensity values were normalized for each embryo to the intensity values at the mid n.c. 14. Embryo 2 is the same from Fig. 4. (**d**) Simulation of the SDD model mimicking the full early development (white continuous line), plotted over the experimental data points from **c**. The simulations were performed to reproduce the experimental trajectories from **c**. Two gradual changes were introduced following build-up of the gradient: an increase in production and a decrease in degradation.

### Protein age as an independent test of models of Bcd gradient formation

Next, we fitted the alternative models to the experimental data from early n.c.14. The SDD model clearly permitted the best general fit to the tFT-Bcd intensity ratio (Fig. 3a), a result that is insensitive to the exact parameters used, as shown in Fig. 1e. Fitting the timer ratio results does not preclude *per se* the *bcd* RNA gradient and the nuclear shuttling models (there are parameter ranges that can fit the ratio), however,only the SDD model can fit both the timer ratio and the intensity spatial profiles. These suggest that long-range Bcd diffusion and Bcd degradation are the primary dynamic processes in forming the Bcd morphogen gradient profile.

The timer provides space-resolved information on the average protein age as well as the gradient shape. We used this dual information to estimate the underlying dynamic parameters assuming a given model, in this case the SDD model. By fitting both the tFT-Bcd ratio and the concentration profiles in early n.c.14 to the SDD model, we estimated the effective Bcd diffusion coefficient and the life-time independently (Fig. 3b, Supplementary Fig. 6). The obtained values for both parameters (D = 3.0 ± 0.7 μm^2^s^-1^ and t_1/2_= 38 ± 7 minutes) are consistent with previous reported measurements. However, a significant advantage here is that both parameters can be estimated from the same embryo at the same moment. The corresponding decay length of the gradient of 82 ± 12 μm is in the range of previously measured values^43^. Including an extended *bcd* RNA source with decay length less than 40µm was also consistent with our data. However, using D ∼ 0.3 μm^2^s^-1^ combined with a long-ranged gradient of *bcd* mRNA - as previously hypothesized^13^ - cannot reproduce the tFT-Bcd ratio, Supplementary Fig. 6n-o. Therefore, our data places a limit on the spatial extent of the *bcd* mRNA gradient. Overall, the tFT-Bcd ratio allows a simultaneous estimation of both diffusion and degradation parameter values and is an independent approach from previous estimations of these parameters.

### The timer can probe degradation mechanisms

Our analysis suggests that Bcd degradation is essential for shaping its concentration gradient. However, it is currently not understood clearly how Bcd is degraded. When assessed in embryonic extracts, Bcd degradation was reported to be proteasome dependent and regulated by the F-box protein Fates-shifted (Fsd) – a specificity factor of the E3 ubiquitin ligase SCF^22^. To further test the proteasome dependency *in vivo*, we injected tFT-Bcd in n.c.14 embryos with the proteasome inhibitor MG132 and imaged them one hour later. We observed that upon proteasomal inhibition there was an increase in the total amount of tFT-Bcd, a steeper gradient profile, and a shift in the timer ratio indicating the presence of older proteins (Fig. 3c-e). However, these effects were not as severe as expected if degradation was completely inhibited, possibly due to partial effect of the drug or the presence of a second degradation pathway. We can fit both DMSO and MG132 injection data with the same parameters using the SDD model (as derived from Fig. 3b), except changing the Bcd lifetime from 38 (DMSO) to 60 (MG132) minutes. To rule out possible indirect effects of the MG132 on the fluorophores, we injected in parallel embryos expressing the construct with the fluorescent tandem timer but without Bcd. This construct is expected to be very stable as it contains no degradation signals. Accordingly, inhibition of the proteasomes in these embryos did not increase the fluorescent ratio (Supplementary Fig. 7a).

We next crossed the timer line into a *fsd*-deficient line (Fig. 3f, Supplementary Fig. 7b). We observe no clear difference between the Fsd mutant and the wt for total amounts of Bcd or gradient profile. The tFT-Bcd ratio is similar between the *fsd* and the wt embryos in n.c.14, Fig 3g, suggesting that Fsd does not play an important role in regulating the mean Bcd lifetime. Our results do not preclude a role for Fsd in Bcd lifetime regulation (e.g. in ensuring robustness), but suggest that Fsd itself is not directly determining the mean Bcd lifetime.

### Protein age during early development

So far we have focused on quantifying the tFT-Bcd ratio in early n.c.14, when the gradient is presumed near equilibrium^10^. In steady-state, the tFT-Bcd ratio is independent of production, and thus gives no information about the Bcd synthesis rate. However, out of steady-state the tFT-Bcd ratio can also be a function of the production rate, see Theory section in Supporting Information.

Thus, we acquired 4D movies of the Bcd timer in developing embryos from the early syncytial blastoderm to gastrulation, Fig. 4a, Supplementary Video 1. Analysis of the sfGFP intensity profile in these embryos matched previous measurements in single-fluorophore tagged lines: a clear Bcd gradient by n.c.10 that increases in intensity until n.c.13 and then starts to decay during n.c.13 and n.c.14 (Fig. 4b)^10,24,31^. The temporal response of the tFT-Bcd ratio suggests that the average Bcd protein age is changing throughout early development (Fig. 4b, lower panel).

To aid interpretation of this new information, we performed *in silico* experiments on the SDD model to distinguish two alternative clearance mechanisms: production cessation or degradation increase. The concentration profiles during the clearance are largely indistinguishable (Fig. 5a insets), but the average protein age increases in the first scenario, and decreases in the second, Fig. 5a. We next allowed the system to reach steady-state, and then changed the production and the degradation rates in a step-wise manner, Fig. 5b. To visualize the timer read-out (age information) and the total intensity (concentration information) simultaneously, we created a phase diagram comparing temporal changes in protein age with the total concentration, Fig. 5b. To reduce the space dimension, we considered only the mean Bcd intensity in the anterior region (other regions are qualitatively similar, Supplementary Fig. 8). Different scenarios for temporal changes in production and degradation resulted in qualitatively distinct trajectories in the phase space (each scenario falls into a different quadrant), potentially enabling them to be distinguished experimentally.

### Synthesis-degradation dynamics during early development

We plotted the experimental phase trajectories (based on Fig. 4a), comparing the tFT-Bcd ratio and the total intensity changes, Fig. 5c, and found all embryos analyzed displayed similar trajectories. Due to the sfGFP folding time (∼ 20 minutes) our results likely represent a delayed readout of around 20-30 minutes in the intensity axis. Comparing these trajectories with Fig. 5b, we find two phases: the establishment of the gradient; and the clearance. The existence of these phases was expected from simple observation of the gradient but our analysis enables dissection of the underlying changes in the production and degradation rates. These phases correlated strongly to the division cycles (Fig. 5c, data point color), suggesting that the beginning of the clearance is likely regulated together with the onset of cellularization. During clearance, the tFT-Bcd ratio increased, consistent with cessation of Bcd synthesis (Fig. 5c and compare trajectories 5 and 6 in Fig. 5b). However, this result does not also preclude a change in Bcd degradation rate. It is noteworthy that we do not observe a full clearance of Bcd during this stage, with some signal remaining until stage 9 in development (Supplementary Fig. 9).

To support this interpretation of the experimental trajectories, we performed a simulation keeping the SDD model structure, but introducing the changes in the production and the degradation rates deduced above (Fig. 5d). More precisely, we started the simulation by building the gradient and then we decreased both the production and the degradation. As shown in Fig. 5d, the simulation results reflect the experimental trajectories closely. This agreement was kept when comparing different regions of the embryo, Supplementary Fig. 8. To achieve this close resemblance, during the second phase, the decrease in the degradation rate has to happen more abruptly than the decrease in the production rate, Supplementary Fig. 10. We tested the effect of impeding diffusion during the clearance phase (i.e. due to cellularization) but no appreciable effect on the phase space was observed. Thus the SDD model structure, combined with time-dependent parameters, can explain the formation of the gradient during early development.

## Discussion

How does the Bcd gradient form? Previously, models of Bcd gradient formation were difficult to distinguish experimentally as they all have similar concentration profiles in early n.c.14 and measurements of dynamic parameters were not sufficiently clear. We have shown here that by measuring the average protein age we can clearly distinguish these models. Our timer analysis revealed the following dynamic and static contraints underlying Bcd gradient formation: (1) Anterior localized Bcd synthesis, with any *bcd* mRNA gradient restricted to < 50μm; (2) Average Bcd protein diffusion around 2-4 μm^2^s^-1^; (3) Spatially homogeneous Bcd degradation with a lifetime of about 40 minutes during early n.c.14; (4) Proteasome (but not Fsd) mediated Bcd degradation; (5) Clearance due to dramatically reduced Bcd synthesis during n.c.14. Our results are most broadly compatible with the SDD model, though the synthesis and degradation parameters are time dependent (as previously suggested^31^). Additionally, we are able to measure the entire extent of the Bcd gradient using MuVi-SPIM. We observe Bcd protein in the posterior pole, a region where mRNA is yet to be observed. This further supports our result that an mRNA gradient plays only a limited role in Bcd protein gradient formation. While the tFT-Bcd reporter has helped illuminate some of the underlying dynamics, there are still unanswered questions including how the dynamic parameters are temporally coordinated and what factors regulate these processes?

More generally, the tFT reporter enables exploration of morphogen dynamics and gradient formation across multiple temporal and spatial scales. Specific advantages include: (1) the analysis is independent of morphogen concentration (though it is sensitive to temporal changes in production); (2) the tFT reporter decouples degradation and diffusion in model fitting; (3) image analysis is straightforward (after careful background subtraction) as the tFT ratio is largely independent of imaging inhomogeneity; and (4) data is extracted from a single time point without need for extended live imaging (and thus avoiding imaging artifacts such as photobleaching). Furthermore, the tFT reporter is sufficiently sensitive to probe degradation regulation. It is becoming increasingly apparent that understanding of developmental patterning requires access to both spatial and temporal information^44,45^. However, accessing this information in the same experiment is challenging. The tFT reporter is a powerful tool for analyzing such systems as it provides both long-ranged spatial information combined with dynamic information.

## Online Methods

### Plasmids and fly lines

All plasmids used in this study were cloned by standard methods. For Bcd constructs all the 3’ and 5’ regulatory regions were conserved. The fluorescent protein fmCherry was evolved in the lab and has the following aminoacidic changes in comparison to mCherry: K52R, K97N, K126R, K143C, K144R, S152T, Y156H, E158G, N201D,T207L, I215V, D232G. The lines 19.C5 (mCherry-sfGFP-Bcd) and the control 79.fd (mCherry-sfGFP) were done by p-element transformation and were later mapped to the crosmosome X and III respectively by crossing them with different balancers. All the other lines were generated by landing-site transgenesis to the chromosome II on a fly strain with background y[1] M{vas-int.Dm}ZH-2A w[*];M{3xP3-RFP/attP’}ZH51C. The wt y/w, and the Bcd^E1^ and fsd^KG02393^ mutant flies were obtained from the Bloomington *Drosophila* Stock Center (stock numbers 42235, 1334 and 12983), and were later crossed with the line 19.C5. The primers used for checking the FSD mutation are: (for) ggcacttgaacagagttacca, (wt specific rev)ggtgaggtaaatttgcactgc and (mutant specific rev) aacaggacctaacgcacagt. Fly stocks were maintained at room temperature on standard agar-cornmeal-yeast media.

### Western Blot

For the collection of the embryos, flies of the corresponding lines were caged 2 to 3 days before the embryos collection. Three 45 min pre-laids were performed, followed by 1 hour lays. “ Stage 4” embryos plates were left at RT for 70 min, and “ Stage 5” embryos plates for 140 min. Then they were collected and snap-frozen in liquid nitrogen. Protein extraction was carried out in High-Urea buffer to a final concentration of 1 embryo/μ l. The following antibodies were used: anti Bicoid (rabbit polyclonal, Santa Cruz Biotechnology), anti GFP (rabbit polyclonal, ab6556; Abcam, Cam- bridge, UK).

### Imaging of live embryos

Images were taken using a custom-made light-sheet microscope^34^, using confocal mode^40^ with a slit size of 30 pixels. In all cases two stacks were acquired simultaneously from two opposing cameras, and then two more stacks were taken after a 90º rotation. The four images obtained were then fused to a single image^34^. The illumination of the sample was performed simultaneously from both sides with 10x objectives, and the images were acquired with 25x 1.1NA water-dipping objectives. With this set up, the full embryo could fit inside the field of view, and a full stack of the embryo, with 201 z-slices separated by 1µm, was acquired typically in 30 sec (exposure time 100-150 msec). See Supplementary Fig. 4 for demonstration of microscope sensitivity and noise (observed fluctuations in Bcd signal were comparable with those previously reported^46^) and uniformity of illumination across the field of view.

For single time points analysis, the images were collected 5-10 minutes after mitotic division 13, when the Bcd gradient is relatively stable^10,31^. Before comparing the ratio of the sfGFP and mCherry signals we removed autofluorescence using linear unmixing^47,48^. In brief, the images were corrected for autofluorescence and background by the subtraction of a weighted *correction* image, taken for this purpose. These images were collected in every experiment as a third channel, using a 488 nm laser and a 594 long-pass filter. For time course analysis, the embryos were collected in 30 min to one-hour long lays, dechorionated approximately 30 min later. Then they were taken to the microscope room, at 18°C, and imaging started approximately 15 min afterwards. The different experiments were performed with a four to six min time resolution, to reduce the effect of photobleaching and phototoxicity. All embryos used in the analysis gastrulated normally.

### Image analysis of the fluorescent timer reporter

To quantify the fluorophore intensities and the tFT-Bcd ratio we performed stereographic projections of the embryo cortical surface^34,49^, Supplementary Fig. 4. In such a 2D projection, image segmentation was straightforward using Ilastik^50^ and we extracted the Bcd intensity profile across the whole embryo. We confirmed the projection accuracy by mapping the coordinates back to the three-dimensional data. The embryo images shown (after autofluorescence correction) are made from a z-projection of half the embryos using the brightest-point method in Fiji. Ratiometric images were generated as outlined previously^38^ in Fiji, and were used only for display purposes.

### Embryo injections

For the injections, we caged newly hatched flies and collected embryos 2 to 3 days later. After at least two 45 min pre-lays, we performed 30 min lays, followed by a 30 min incubation at room temperature, and a 40 sec dechorionation in 50% bleach. The embryos were mounted for injections on a coverslip with heptane glue. Treated and control embryos were mounted side-by-side. After a 5 min de-hydration, they were covered with a drop of halocarbon oil 700/27 (1:2; Sigma-Aldrich). The injection needles were pulled from borosilicate glass capillaries (1.2-mm outer diameter, 0.94-mm inner diameter; Harvard Apparatus), using a P-97 Flamming/brown puller (Sutter Instrument). The injections were performed using an Eppendorf microinjector model 5242. The injection time was 0.5 sec in every case, and the injection pressure was calibrated for each needle and liquid to produce a drop of the same approximate volume (by simple observation in comparison to the size of a mesh, aiming to a 5% of the embryo’ s volume). The injections were performed in the posterior pole of the embryos. For the proteasome inhibitor MG132 (C2211; Sigma-Aldrich) we used a concentration of 1 mM in DMSO. Control embryos were injected with DMSO alone. For the RNA a concentration of 100 ng/μl in RNAses free water was used (controls were injected with water).

More details on the determination of the maturation rates of the fluorophores are available in the Supplementary Information.

### Model fitting

To fit the tFT-ratio we needed to account for the different intensity of the sfGFP and mCherry fluorophores. For this we considered two options: (1) use *a priori* information and normalize the profiles at large distances (as in Fig. 2e inset), where we expected only *old* protein; (2) or leave the intensity normalization factor as a free parameter. The consequence of the latter scenario is to increase the uncertainty range on the diffusion and degradation parameters, Supplementary Fig. 6 and Supplementary Information. All results presented in the main figures uses either un-normalized tFT-ratios or use an additional parameter to account for intensity scaling.

### Parameter estimation

For each model, we calculated 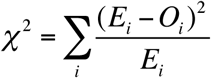, where *E*_*i*_ and *O*_*i*_ represent the model prediction and experimental observation for each position *x*_*i*_. Using alternative fitting, such as a simple least-squares analysis did not significantly alter the main conclusions, data not shown. Error bars on model fits were calculated by calculating the parameter range for which χ^2^ < 10^−3^. Fits were performed using different approximations for FRET effects, Supplementary Fig. 6.

### Simulations and Modeling

Mathematical details of all models considered are described in the Supplementary Material. Briefly, models were numerically solved with Bcd insertion at x=0 μm and no-flux boundary condition at L=500 μm, Supplementary Fig. 1 for further simulation results. All calculations and simulations for Fig. 1-3 were performed in Matlab. The simulations for Fig. 5 were performed in R, using the package Reactdiff. Details of analytical approaches to calculating the protein age are described in the Supplementary Information and Supplementary Fig. 10.

## Acknowledgements

We thank the members of the Knop and Hufnagel laboratories for helpful comments and discussions. In particular, we acknowledge technical help from Gustavo de Medeiros, Bálint Balázs, Marvin Albert and Birgit Besenbeck. We thank the mechanical and electronics workshop of the European Molecular Biology Laboratory (EMBL) for customized hardware. The EMBL Advanced Light Microscopy Facility is acknowledged for support in image acquisition and analysis. This work was supported by the European Molecular Biology Laboratory (L.D., T.S., T.E.S. and L.H.). T.E.S. was further supported by the EMBL Interdisciplinary Postdoc Programme (EIPOD) under Marie Curie Actions COFUND and support from the Kavli Institute for Theoretical Physics, Santa Barbara. LD was supported from a postdoc EcTOP2 fellowhip from the CellNetworks cluster of the excellence initiative of the German research council. L.D., M.K. and L.H. acknowledge support by the CellNetworks, University of Heidelberg, Germany in the context of the EcTop2 project.

## Author contributions

L.D, M.K., T.E.S. and L.H. designed the study, discussed results and wrote the manuscript. L.D., D.K., T.S., S.R. and T.E.S. carried out experiments; L.D., I.K., and T.E.S. analyzed the data.

## References

1. Wolpert, L. Positional information and the spatial pattern of cellular differentiation. J. Theor. Biol. 25, 1–47 (1969).

2. Lander, A. D. Pattern, growth, and control. Cell 144, 955–969 (2011).

3. Ribes, V. & Briscoe, J. Establishing and Interpreting Graded Sonic Hedgehog Signaling during Vertebrate Neural Tube Patterning: The Role of Negative Feedback. CSH Persp. Biol. 1, a002014–a002014 (2009).

4. Rogers, K. W. & Schier, A. F. Morphogen gradients: from generation to interpretation. Annu. Rev. Cell Dev. Biol. 27, 377–407 (2011).

5. Muller, P., Rogers, K. W., Yu, S. R., Brand, M. & Schier, A. F. Morphogen transport. Development 140, 1621–1638 (2013).

6. Driever, W. & Nüsslein-Volhard, C. A gradient of bicoid protein in Drosophila embryos. Cell 54, 83–93 (1988).

7. Driever, W. & Nüsslein-Volhard, C. The bicoid protein determines position in the Drosophila embryo in a concentration-dependent manner. Cell 54, 95–104 (1988).

8. Frigerio, G., Burri, M., Bopp, D., Baumgartner, S. & Noll, M. Structure of the segmentation gene paired and the Drosophila PRD gene set as part of a gene network. Cell 47, 735–746 (1986).

9. Berleth, T. et al. The role of localization of bicoid RNA in organizing the anterior pattern of the Drosophila embryo. EMBO J. 7, 1749–1756 (1988).

10. Gregor, T., Wieschaus, E. F., McGregor, A. P., Bialek, W. & Tank, D. W. Stability and Nuclear Dynamics of the Bicoid Morphogen Gradient. Cell 130, 141–152 (2007).

11. Houchmandzadeh, B., Wieschaus, E. & Leibler, S. Establishment of developmental precision and proportions in the early Drosophila embryo. Nature 415, 798–802 (2002).

12. Coppey, M., Berezhkovskii, A. M., Kim, Y., Boettiger, A. N. & Shvartsman, S. Y. Modeling the bicoid gradient: Diffusion and reversible nuclear trapping of a stable protein. Dev. Biol. 312, 623–630 (2007).

13. Spirov, A. et al. Formation of the bicoid morphogen gradient: an mRNA gradient dictates the protein gradient. Development 136, 605–614 (2009).

14. Hecht, I., Rappel, W.-J. & Levine, H. Determining the scale of the Bicoid morphogen gradient. Proc. Natl. Acad. Sci. U.S.A. 106, 1710–1715 (2009).

15. Dilão, R. & Muraro, D. mRNA diffusion explains protein gradients in Drosophila early development. J. Theor. Biol. 264, 847–853 (2010).

16. Kavousanakis, M. E., Kanodia, J. S., Kim, Y., Kevrekidis, I. G. & Shvartsman, S. Y. A compartmental model for the bicoid gradient. Developmental Biology. Dev. Biol. 345, 12–17 (2010).

17. Grimm, O., Coppey, M. & Wieschaus, E. Modelling the Bicoid gradient. Development 137, 2253–2264 (2010).

18. Abu-Arish, A., Porcher, A., Czerwonka, A., Dostatni, N. & Fradin, C. High mobility of bicoid captured by fluorescence correlation spectroscopy: implication for the rapid establishment of its gradient. Biophys. J. 99, L33–35 (2010).

19. Castle, B. T., Howard, S. A. & Odde, D. J. Assessment of Transport Mechanisms Underlying the Bicoid Morphogen Gradient. Cel. Mol. Bioeng. 4, 116–121 (2011).

20. Sigaut, L., Pearson, J. E., Colman-Lerner, A. & Ponce Dawson, S. Messages Do Diffuse Faster than Messengers: Reconciling Disparate Estimates of the Morphogen Bicoid Diffusion Coefficient. PLoS Comput. Biol. 10, e1003629 (2014).

21. Drocco, J. A., Grimm, O., Tank, D. W. & Wieschaus, E. Measurement and perturbation of morphogen lifetime: effects on gradient shape. Biophys. J. 101, 1807–1815 (2011).

22. Liu, J. & Ma, J. Fates-shifted is an F-box protein that targets Bicoid for degradation and regulates developmental fate determination in Drosophila embryos. Nat. Cell. Biol. 13, 22–29 (2011).

23. Liu, J., He, F. & Ma, J. Morphogen gradient formation and action. Fly 5, 242–246 (2011).

24. Bergmann, S. et al. Pre-Steady-State Decoding of the Bicoid Morphogen Gradient. PLoS Biol. 5, e46 (2007).

25. Bergmann, S., Tamari, Z., Schejter, E., Shilo, B.-Z. & Barkai, N. Re-examining the Stability of the Bicoid Morphogen Gradient. Cell 132, 15–17 (2008).

26. Bialek, W., Gregor, T., Tank, D. W. & Wieschaus, E. F. Response: Can We Fit All of the Data? Cell 132, 17–18 (2008).

27. Saunders, T. & Howard, M. When it pays to rush: interpreting morphogen gradients prior to steady-state. Phys. Biol. 6, 046020 (2009).

28. de Lachapelle, A. M. & Bergmann, S. Precision and scaling in morphogen gradient read-out. Mol. Sys. Biol. 6, 351 (2010).

29. Jaeger, J. A matter of timing and precision. Mol. Sys. Biol. 6, 427 (2010).

30. de Lachapelle, A. M. & Bergmann, S. Pre-steady and stable morphogen gradients: can they coexist? Mol. Sys. Biol. 6, 428 (2010).

31. Little, S. C., Tkačik, G., Kneeland, T. B., Wieschaus, E. F. & Gregor, T. The Formation of the Bicoid Morphogen Gradient Requires Protein Movement from Anteriorly Localized mRNA. PLoS Biol. 9, e1000596 (2011).

32. Wieschaus, E. in Essays on Developmental Biology, Part B 117, 567–579 (Elsevier, 2016).

33. Khmelinskii, A. et al. Tandem fluorescent protein timers for in vivo analysis of protein dynamics. Nat. Biotechnol. 30, 708–714 (2012).

34. Krzic, U., Gunther, S., Saunders, T. E., Streichan, S. J. & Hufnagel, L. Multiview light-sheet microscope for rapid in toto imaging. Nat. Meth. 9, 730–733 (2012).

35. Tomer, R., Khairy, K., Amat, F. & Keller, P. J. Quantitative high-speed imaging of entire developing embryos with simultaneous multiview light-sheet microscopy. Nat. Meth. 9, 755–763 (2012).

36. Dalessi, S., Neves, A. & Bergmann, S. Modeling morphogen gradient formation from arbitrary realistically shaped sources. J. Theor. Biol. 294, 130–138 (2012).

37. Pédelacq, J.-D., Cabantous, S., Tran, T., Terwilliger, T. C. & Waldo, G. S. Engineering and characterization of a superfolder green fluorescent protein. Nat. Biotechnol. 24, 79–88 (2005).

38. Khmelinskii, A. & Knop, M. in Computational systems biology (eds. Ireton, R., Montgomery, K., Bumgarner, R., Samudrala, R. & McDermott, J.) 1174, 195–210 (Springer New York, 2014).

39. Baumgart, E. & Kubitscheck, U. Scanned light sheet microscopy with confocal slit detection. Opt. Express 20, 21805–21814 (2012).

40. de Medeiros, G. et al. Confocal multiview light-sheet microscopy. Nat. Comm. 6, 1–8 (2015).

41. Stelzer, E. H. K. Light-sheet fluorescence microscopy for quantitative biology. Nat. Meth. 12, 23–26 (2015).

42. Khmelinskii, A. et al. Incomplete proteasomal degradation of green fluorescent proteins in the context of tandem fluorescent protein timers. Mol. Biol. Cell 27, 360–370 (2016).

43. Liu, F., Morrison, A. H. & Gregor, T. Dynamic interpretation of maternal inputs by the Drosophila segmentation gene network. Proc. Natl. Acad. Sci. U.S.A. 110, 6724–6729 (2013).

44. Alexandre, C., Baena-Lopez, A. & Vincent, J.-P. Patterning and growth control by membrane-tethered Wingless. Nature 505, 180–185 (2014).

45. Kicheva, A. et al. Coordination of progenitor specification and growth in mouse and chick spinal cord. Science 345, 1254927 (2014).

46. Gregor, T., Tank, D. W., Wieschaus, E. F. & Bialek, W. Probing the Limits to Positional Information. Cell 130, 153–164 (2007).

47. Dickinson, M. E., Bearman, G., Tille, S., Lansford, R. & Fraser, S. E. Multi-spectral imaging and linear unmixing add a whole new dimension to laser scanning fluorescence microscopy. BioTechniques 31, 1272–1278 (2001).

48. Kraus, B., Ziegler, M. & Wolff, H. Linear fluorescence unmixing in cell biological research. Mod. Res. Educ. Top. Microscopy 863–872 (2007).

49. Schmid, B. et al. High-speed panoramic light-sheet microscopy reveals global endodermal cell dynamics. Nat. Comm. 4, 1–10 (2013).

50. Sommer, C., Straehle, C. & Kothe, U. ilastik: Interactive learning and segmentation toolkit. IEEE Int. Symp. Biomed. Imaging (2011).

